# Fitting photosynthetic carbon dioxide and irradiance response curves for C_4_ leaves

**DOI:** 10.1101/2025.06.03.657559

**Authors:** Russell Woodford, Maria Ermakova, Robert T. Furbank, Susanne von Caemmerer

## Abstract

Gas exchange measurements provide crucial insights into the complex mechanisms of photosynthesis. Responses of CO_2_ assimilation rate (*A*) to intercellular CO_2_ partial pressure (*C*_i_) and irradiance (*I*) link gas exchange measurements to the underlying photosynthetic biochemistry of a leaf. The unique biochemistry and leaf anatomy which distinguish C_4_ photosynthesis make it necessary to apply models and fitting routines which appropriately parameterise and incorporate these characteristics. Here we provide updates to the C_4_ photosynthesis model by improving the parameterisation of cyclic electron flow in C_4_ photosynthesis using experimentally derived values from *S. viridis.* We additionally describe two fitting routines for assessing C_4_ photosynthesis based on the updated model. Fitting of a CO_2_ response curve (*A/C*_i_) provides estimates of maximum PEP carboxylase activity (*V*_pmax_) and maximum Rubisco activity (*V*_cmax_), and calculates the electron transport rate (*J*) needed to sustain the measured CO_2_ assimilation rate (*A*). Fitting of an irradiance response curve (*A/I*) provides estimates for the maximum electron transport rate (*J*_max_), day respiration rate (*R*_d_), the quantum yield (*ϕ*_CO2_) and light compensation point (*Γ*_light_). Values of the above output parameters are provided at both the measurement temperature and at 25 °C for ease of comparative reporting. The fitting tool has been designed in Microsoft Excel to minimise barriers to entry and enable simplicity of fitting while simultaneously catering to individuals with diverse expertise and experience in C_4_ gas exchange modelling.

## Introduction

CO_2_ assimilation by leaves depends on the response of underlying photosynthetic biochemistry to environmental variables such as light, CO_2_ and temperature. The Farquhar et al. model of C_3_ photosynthesis has associated the gas exchange of C_3_ leaves with their fundamental biochemical mechanisms (Farquhar *et al*., 1980; von Caemmerer and Farquhar, 1981). Gas exchange measurements of CO_2_ response curves are now routinely used to extract biochemical parameters such as the maximum activity (*V*_cmax_) of ribulose-1,5-bisphosphate carboxylase/oxygenase (Rubisco), maximum rate of electron transport for the given light intensity (*J*), and triose phosphate limitation (*TPU*) from C_3_ leaves at high CO_2_ partial pressures using an easy curve fitting routine (Sharkey *et al*., 2007). Here we have developed a curve fitting routine for CO_2_ response and light response curves of the C_4_ photosynthetic pathway based on the updated model of C_4_ photosynthesis (von Caemmerer, 2021). We have also updated values for the fraction of cyclic electron flow (*f*_cyc_) and number of protons shuttled per cyclic electron flow (*H*_Jcyc_) in C_4_ photosynthesis.

The C_4_ photosynthetic pathway is complex and involves a biochemical CO_2_ concentration mechanism (CCM) operating across mesophyll and bundle sheath cells. In the mesophyll, CO_2_ is initially fixed by phosphoenolpyruvate (PEP) carboxylase (PEPC) into C_4_ acids, which are then decarboxylated in the bundle sheath to supply CO_2_ for Rubisco. The C_4_ CCM allows Rubisco to operate at high CO_2_ partial pressures, overcoming its low affinity for CO_2_, largely inhibiting its oxygenation reaction and reducing photorespiration rates. Both the structure of the bundle sheath cell wall (which has a low permeability to CO_2_) and the relative biochemical capacities of the C_3_ cycle in the bundle sheath and C_4_ acid cycle operating across the mesophyll–bundle sheath interface contribute to the high CO_2_ partial pressure in the bundle sheath. This leads to distinct CO_2_ response (*A*/*C*_i_) curves of the C_4_ photosynthetic pathway, characterised by a steep initial slope and a saturation of CO_2_ assimilation rate at lower *C*_i_, in comparison to *A*/*C*_i_ curves of C_3_ species (von Caemmerer, 2021; von Caemmerer *et al*., 2012). Furthermore, due to the C_4_ CCM, CO_2_ compensation points are low and independent of O_2_ partial pressures. C_4_ light response (*A*/*I*) curves can be characterised by a lack of saturation, even at irradiance above 2000 µmol m^-2^ s^-1^ (Ermakova *et al*., 2019; Leakey *et al*., 2006), and a quantum yield independent of temperature (Ehleringer and Björkman, 1977).

Although C_4_ plants are not limited by CO_2_ availability under ambient conditions, the electron transport reactions producing ATP and NADPH could limit the rate of C_4_ photosynthesis at high irradiance (Ermakova *et al*., 2019; von Caemmerer and Furbank, 2016). Two major electron transfer pathways differentially contribute to ATP and NADPH production: linear electron flow (*J*) and cyclic electron flow (*J*_cyc_). *J* involves Photosystem II (PSII), Cytochrome *b*_6_*f* (Cyt*b*_6_*f*), Photosystem I (PSI), ferredoxin:NADP^+^ oxidoreductase (FNR) and ATP synthase. This produces ATP and NADPH in an ATP: NADPH ratio of ∼2.6:2 (Kramer and Evans, 2011). However, for each molecule of CO_2_ fixed, C_4_ photosynthesis requires an ATP: NADPH ratio of at least 5:2 (von Caemmerer and Furbank, 2016). *J*_cyc_ is an alternative pathway of electron transport used to increase ATP yield (Kramer and Evans, 2011; Yamori and Shikanai, 2016). In *J*_cyc_, electrons from PSI bypass FNR and return to Cyt*b*_6_*f* via either a chloroplast NADPH dehydrogenase-like (NDH) complex–dependent pathway or a Proton Gradient Regulation 5 (PGR5)–dependent pathway, leading to the production of ATP but not NADPH (Yamori and Shikanai, 2016). To accommodate their increased ATP requirements, it has been proposed that C_4_ species upregulate *J*_cyc_ (Ishikawa *et al*., 2016; Munekage and Taniguchi, 2016; Nakamura *et al*., 2013). However, previous iterations of the C_4_ photosynthesis model have underestimated the contributions of *J*_cyc_ to total electron transport, potentially miscalculating the influence *J*_cyc_ could have on gas exchange measurements, particularly when fitting *A*/*I* curves. To improve the accuracy and applicability of the model, better parameterisation of the fraction of cyclic electron flow (*f*_cyc_) and the number of protons pumped per *J*_cyc_ (*H*_Jcyc_) has been implemented.

### The C_4_ photosynthetic model with additional detail of the electron transport rate

The first models to capture the C_4_ photosynthetic biochemistry were designed by Berry and Farquhar (1978) and Peisker (1979). The Berry and Farquhar model did not provide analytical solutions but was able to predict high bundle sheath CO_2_ partial pressures and their dependence on bundle sheath conductance. Many of the gas exchange characteristics of C_4_ photosynthesis observed with intact leaves could be predicted by these models. Von Caemmerer and Furbank (1999) revised and expanded these original models with analytical solutions. Most recently, von Caemmerer (2021) again revised and updated this model with Rubisco and PEP carboxylase kinetic parameters derived from *Setaria viridis* and incorporated expanded electron transport functions and temperature functions, improving the overall accuracy of the model.

Here we have updated the model of von Caemmerer (2021) with a better parameterised cyclic electron flow by assigning updated values to the number of protons pumped per electron in *J*_cyc_ (*H*_Jcyc_) and the fraction of total electron transport that can be attributed to *J*_cyc_ (*f*_cyc_). This is achieved through both a theoretical and empirical approach. As *J* is proposed to produce ATP and NADPH in a ratio of ∼2.6:2 (=1.3) while *J*_cyc_ is proposed to only produce ATP, the *f*_cyc_ needed to fulfil the minimum ATP: NADPH ratio of 5:2 (=2.5) required for C_4_ photosynthesis can be theoretically estimated for varying *H*_Jcyc_ using Equations 1-3.

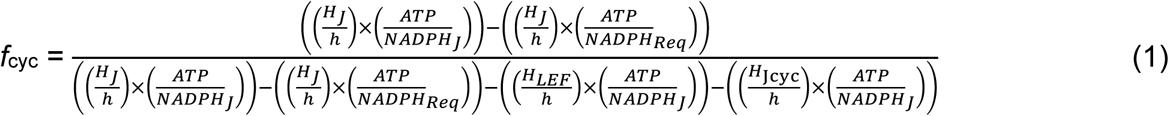

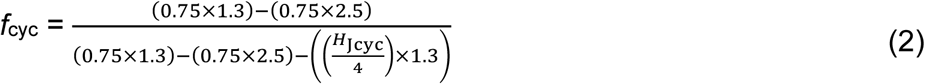

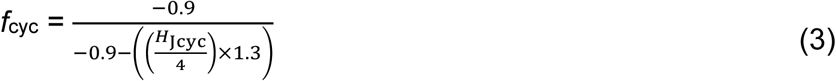

Where:

*H*_J_ = proton: electron ratio of linear electron flow = 3;

*H*_Jcyc_ = proton: electron ratio of cyclic electron flow;

*h* = proton: ATP production ratio of ATP synthase = 4;

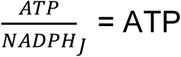: NADPH ratio produced by linear electron flow = 1.3;

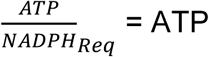: NADPH ratio required by C_4_ photosynthesis = 2.5.

This allows for the range of *f*_cyc_ values required to fulfill the minimum ATP requirements of C_4_ photosynthesis to be calculated, providing a minimum *f*_cyc_ of 0.41 when *H*_Jcyc_ *=* 4, and maximum *f*_cyc_ of 0.58 when *H*_Jcyc_ *=* 2 (Figure 1). It is proposed that the NDH-dependent pathway provides the dominant contribution to *J*_cyc_ in C_4_ photosynthesis, as opposed to the PGR5-dependent pathway which has yet to be shown to contribute to ATP synthesis (Ermakova *et al*., 2024; Ogawa *et al*., 2023; Woodford *et al*., 2024). As the NDH-dependent pathway is proposed to move 4H^+^ per electron (Strand *et al*., 2017), this suggests the value of *f*_cyc_ is likely closer to the lower predicted value.

**Figure 1.**
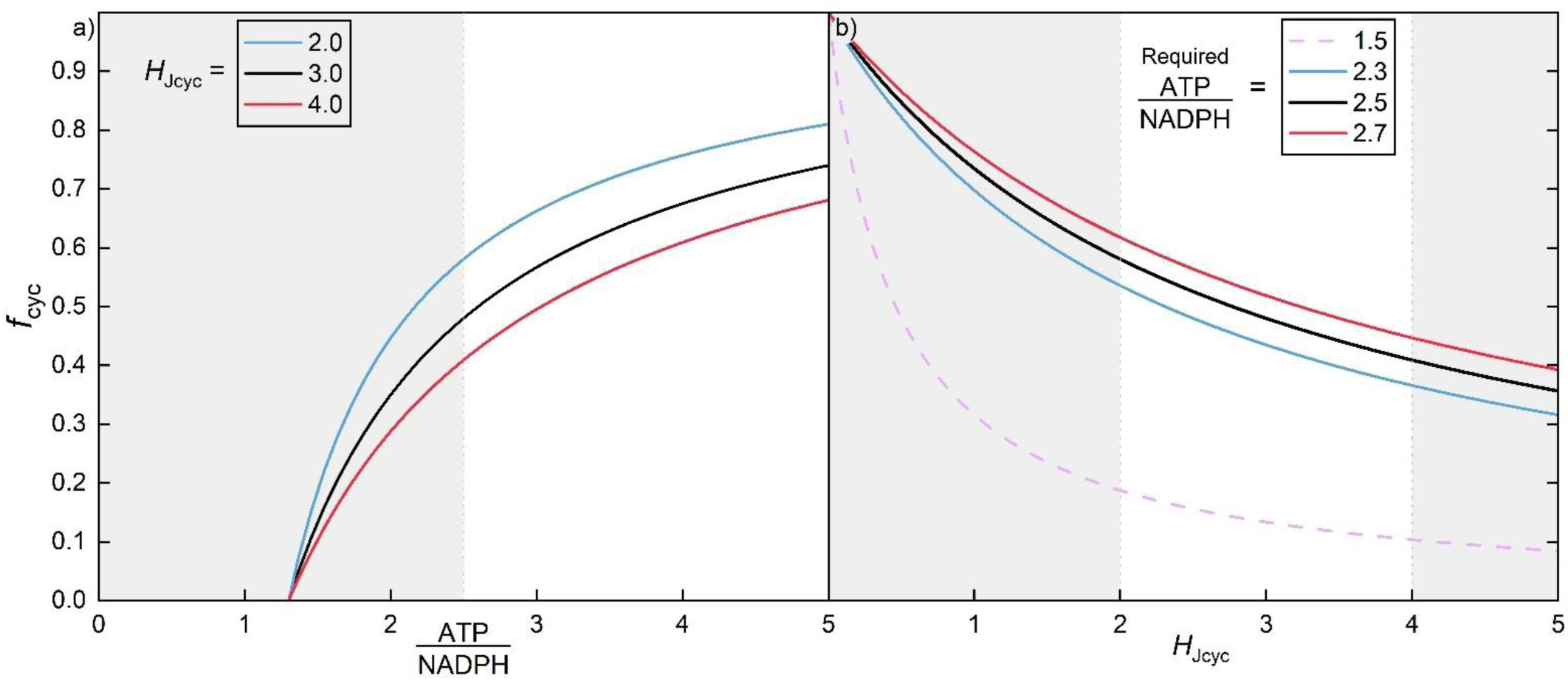
Dependence of the fraction of electron transport occurring through cyclic electron flow (*f*_cyc_) on the ATP: NADPH requirement of C_4_ photosynthesis and the number of protons per electron pumped by cyclic electron flow (*H*_Jcyc_). (**a**) The *f*_cyc_ required to accommodate the minimal ATP: NADPH requirement of C_4_ photosynthesis (values above 2.5, white range) is dependent on *H*_Jcyc_. Increased *f*_cyc_ is required to accommodate C_4_ photosynthesis when fewer protons are pumped per electron. (**b**) The range of *f*_cyc_ values required to accommodate varying ATP: NADPH across physiologically reasonable *H*_Jcyc_ (values between 2 and 4, white range). Lower *f*_cyc_ is required to accommodate C_4_ photosynthesis when a higher number of protons per electron are pumped. The theoretical *f*_cyc_ required for C_3_ photosynthesis (ATP: NADPH = 1.5) is less than half that for C_4_ photosynthesis.

Values of *f*_cyc_ within this theoretical range are also supported empirically. Treatment of *S. viridis* bundle sheath cells with methyl viologen, an electron acceptor from PSI competing with *J*_cyc_, demonstrates that *f*_cyc_ is 0.84 in bundle sheath cells (Ermakova *et al*., 2024). In C_3_ plants, which only have mesophyll cells and little NDH in optimal conditions, *f*_cyc_ is estimated to be 0.12 (Avenson *et al*., 2005). Due to similarities of their electron transport chains, *f*_cyc_ in C_3_ and C_4_ mesophyll cells could be assumed similar (Munekage, 2016). However, analysis of the light-driven proton flux across the thylakoid membrane (representing *J*_cyc_ and *J*) and the relative PSII electron transport rate (representing *J*) in PGR5-deficient *S. viridis* suggest that PGR5-mediated *J*_cyc_ accounts for at least 20% of mesophyll electron transport (*J*_m_) (Woodford *et al*., 2024). As about 40% of total leaf chlorophyll is localised to bundle sheath cells in *S. viridis* (Ermakova *et al*., 2021a), the leaf-level *f*_cyc_ can be calculated using Equation 4.

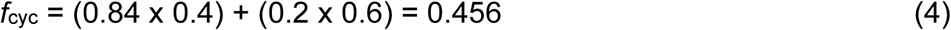

To fulfill the ATP: NADPH requirements of C_4_ photosynthesis, this would require *H*_Jcyc_ ∼3.3, thereby necessitating a 65% contribution of NDH and 35% contribution of PGR5 to total leaf *J*_cyc_ (Equation 5).

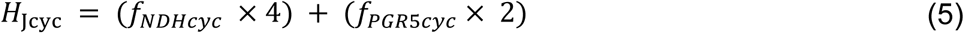

Previous empirical values for *f*_cyc_ determined from measurements of the quantum efficiency of oxygen evolution under limiting light align with these calculations, with *f*_cyc_ values ranging from 0.32 to 0.62, and averaging 0.45 (Bjorkman and Demmig, 1987; Yin and Struik, 2012). Estimates of *f*_cyc_ calculated using measurements of P700^+^ absorbance, chlorophyll fluorescence and oxygen evolution light response curves, provided an *f*_cyc_ value of 0.53 for NADP-ME species (Kiirats *et al*., 2010; Yin and Struik, 2012). Given the strong consistency in these values, we have therefore assigned a *H*_Jcyc_ of 3.4, and a *f*_cyc_ value of 0.45. However, it is worth acknowledging that *f*_cyc_ could be higher to accommodate CO_2_ leakiness of bundle sheath cells, H^+^ leakage of thylakoid membranes or additional, unaccounted for ATP requirements, particularly during photosynthetic induction, under non-steady-state conditions or at high temperatures (Bukhov *et al*., 1999; Ermakova *et al*., 2023; Groth and Junge, 1993; Wang *et al*., 2022).

### C_4_ photosynthesis fitting routines

Using the von Caemmerer (2021) model and incorporating the improvements detailed above, we have developed an Excel fitting routine for *A*/*C*_i_ and *A*/*I* curves measured on C_4_ leaves. The fitting routines are based on the Equations 21 and 40, and Table 1 from von Caemmerer (2021). The essential parameters used in the fitting routine are presented here in Table 1.

**Table 1.**
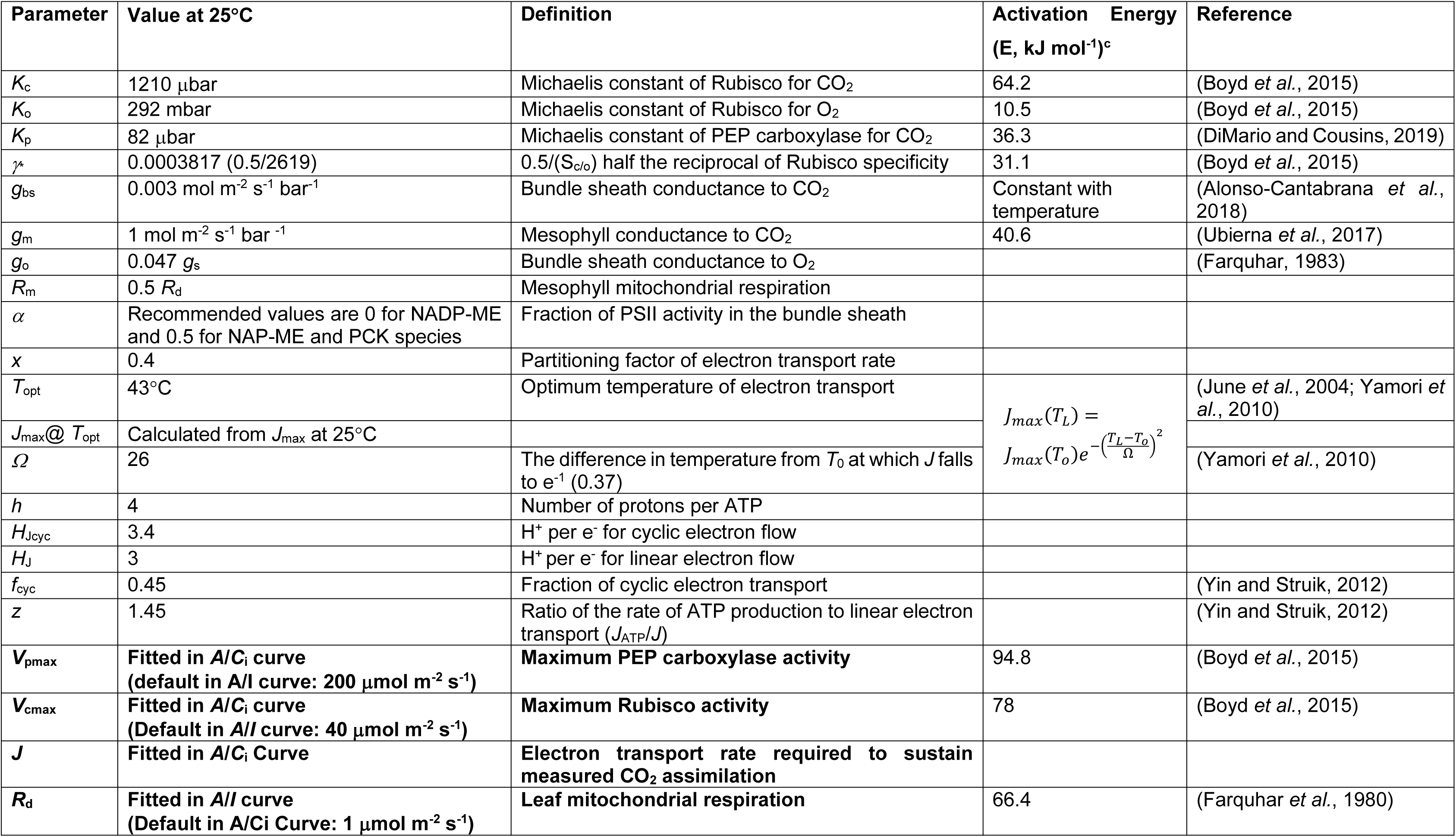
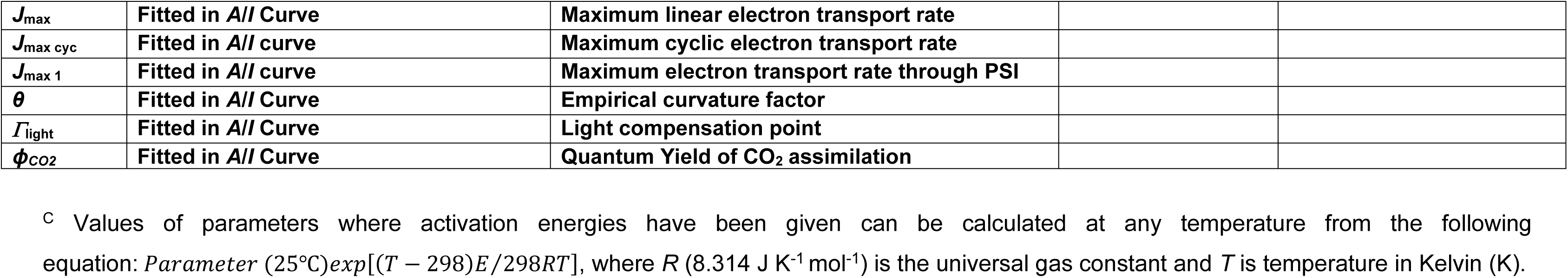
Photosynthetic parameters used in the fitting routines.

### Fitting a CO_2_ response curve

Before fitting an *A*/*C*_i_ curve, one first updates leaf temperature, atmospheric pressure, irradiance and O_2_ concentration in the spreadsheet. Depending of the biochemical subtype of the species examined, the fraction of PSII active in the bundle sheath (*α*) is set to a value between 0 and 1. NADP-ME monocot species such as *Zea mays* and *Sorghum bicolor* have little or no PSII activity in the bundle sheath in optimal conditions, whereas NADP-ME dicots, NAD-ME and PCK species can have high bundle sheath PSII activity (von Caemmerer, 2021 and references therein). The *α* value should be updated to reflect this. CO_2_ and O_2_ concentration are frequently expressed in mole fractions (i.e., µmol mol^-1^, ppm, µL L^-1^) and can be entered as such. These CO_2_ and O_2_ mole fractions are then converted to partial pressure with the use of the given atmospheric pressure, enabling the use of partial pressure for the units of CO_2_ and O_2_ in all calculations. This is justified as the chemical activity of a dissolved gas is proportional to its gas-phase, and thus the partial pressure of a gas existing in equilibrium with solution is a better measure of its chemical activity than dissolved concentrations (Badger and Collatz, 1977; Sharkey *et al*., 2007). Furthermore, the mole fraction of CO_2_ is the same at sea level and at high altitude, but the partial pressure declines with increasing elevation as atmospheric pressure declines.

To fit an *A*/*C*_i_ curve one enters the measured values for CO_2_ assimilation rate (*A*) and intercellular CO_2_ (*C*_i_). In the fitting routine of an *A*/*C*_i_ curve for C_3_ photosynthesis, users must choose which data points are used to calculate the Rubisco limited, electron transport limited or triose phosphate limited rates (Sharkey *et al*., 2007; von Caemmerer and Farquhar, 1981). In the C_4_ photosynthesis model such choices do not need to be made. Simply pressing the calculate button will solve for *V*_cmax_ (the maximum Rubisco activity) and *V*_pmax_ (the maximum PEPC activity) values which minimise the sum of squares between experimental and modelled data. After converging on a solution, the *V*_cmax_ and *V*_pmax_ will be the output for the modelled enzyme-limited CO_2_ assimilation rate, which is appropriate at high irradiance. At the same time, electron transport rate, *J,* required to sustain CO_2_ assimilation rate is also calculated from the enzyme-limited rate at *C*_i_. These values are given at both the measurement temperature and calculated at 25°C for reporting purposes.

### Interpretation of the *A*/*C*_i_ fitting results and resolving limitations

At high irradiance, the C_4_ photosynthetic model predicts that *A* is limited by *V*_pmax_ at low *C*_i_ and co- limited by *V*_cmax_ and *J* at high *C*_i_. Together, *V*_pmax,_ *V*_cmax_ and *J*, given in units of µmol m^-2^ s^-1^, describe C_4_ *A*/*C*_i_ curves. *V*_cmax_ and *V*_pmax_ can be compared to *in vitro* measurements of those activities (Cousins *et al*., 2007) while electron transport rate can be related to the Cytochrome *b*_6_*f* content in leaves and to a spectrophotometric assay for cytochrome *f* (Ermakova *et al*., 2019; Heyno *et al*., 2022; Yamori *et al*., 2010). Leaf nitrogen is another parameter that is a useful corelate with CO_2_ assimilation rate, particularly in field studies (Ghannoum *et al*., 2005).

As Rubisco and electron transport rate co-limit photosynthetic flux at high *C*_i_ and high light, distinguishing these limitations might be challenging. For example, fitting of *A*/*C*_i_ from *S. viridis* plants overexpressing the Rieske FeS protein of Cytochrome *b*_6_*f*, which have increased electron transport capacity but no changes in Rubisco content (Ermakova *et al*., 2019), suggests that both *V*_cmax_ and *J* are increased compared to control (Figure 2). If the primary limitation needs to be resolved, strategies can be employed to help disentangle these parameters. We recommend making a series of *A* measurements at high *C*_i_ with increasing irradiances. If increasing irradiance results in increased CO_2_ assimilation rate, it is likely that electron transport is the limiting step rather than *V*_cmax_. When *J* is limiting, one must be aware that *V*_cmax_ does not represent the maximum Rubisco activity, but rather the activity allowed by the electron transport rate at given irradiance. Measurements of *A*/*I* curves on the same leaves can provide further insight into Rubisco and electron transport limitations (see below).

**Figure 2.**
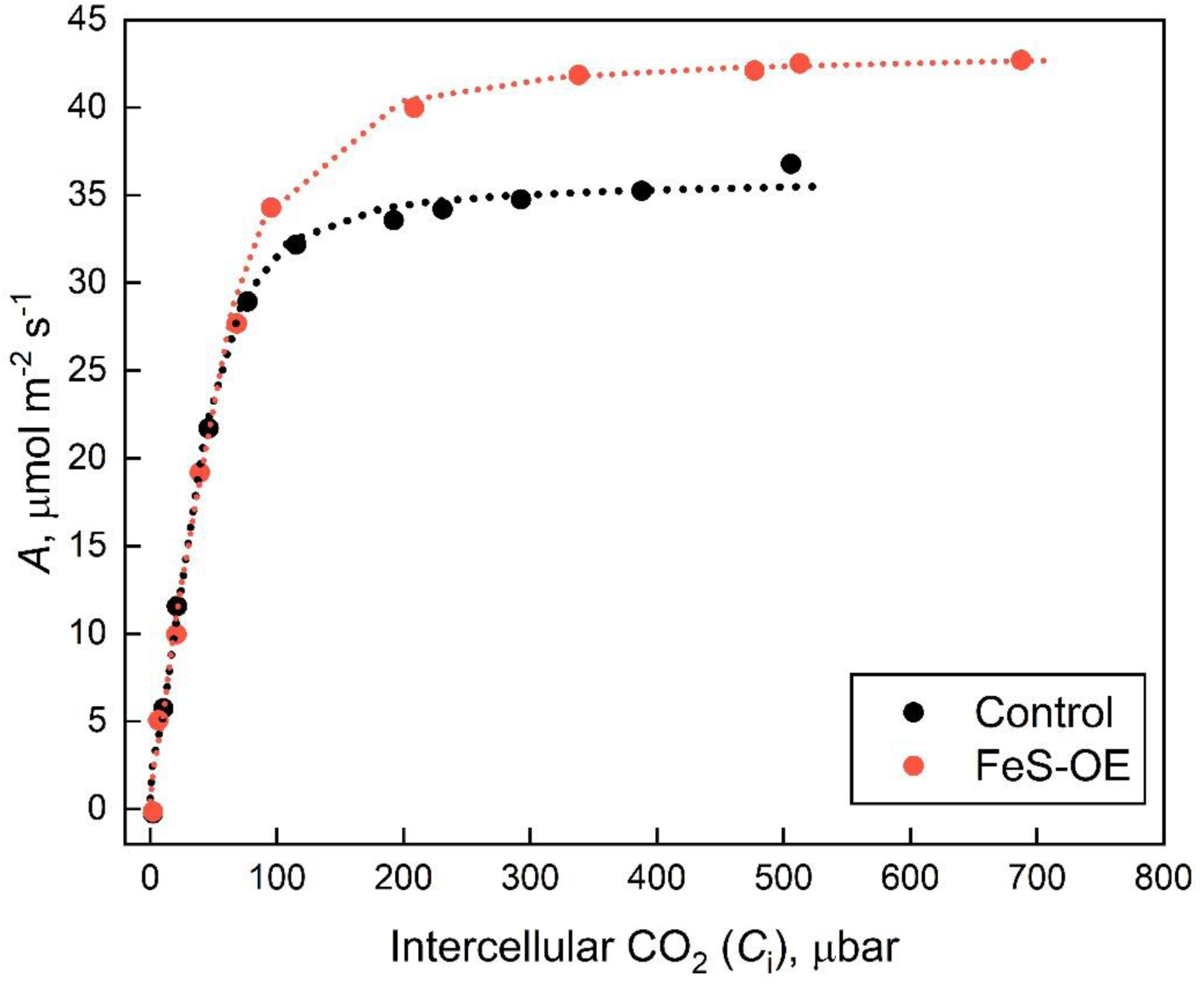
Fitting of the CO_2_ response curves of CO_2_ assimilation (*A*) from Control and Rieske FeS overexpression (FeS-OE) *S. viridis* plants. Data originally reported in Ermakova *et al*. (2019). Dotted lines represent the fit of modelled data to the measured data points. The fitted values for FeS-OE plants are *V*_pmax_ = 152.17 µmol m^-2^ s^-1^, *V*_cmax_ = 47.81 µmol m^-2^ s^-1^ and *J* = 166.83 µmol m^-2^ s^-1^. For WT, *V*_pmax_ = 179.16 µmol m^-2^ s^-1^, *V*_cmax_ = 39.36 µmol m^-2^ s^-1^ and *J* = 141.26 µmol m^-2^ s^-1^.

### Important *A*/*C*_i_ fitting considerations

It is thought that in most C_4_ leaves, carbonic anhydrase activity is sufficiently high and does not limit photosynthetic flux (Cousins *et al*., 2008; Hatch and Burnell, 1990). As such, we have not included carbonic anhydrase activity as a limitation here. We also do not recommend that bundle sheath conductance is fitted using our routine. The pre-set value of *g*_bs_ = 0.003 mol m^-2^ s^-1^ bar^-1^ is consistent with measured values (Table 1, Bellasio *et al*., 2024; Danila *et al*., 2021). The value needs to be low to reproduce the observed lack of oxygen sensitivity of C_4_ photosynthesis (von Caemmerer, 2000, Fig. 4.6).

Mesophyll conductance (*g*_m_), the ease of CO_2_ diffusion from the intercellular airspace to the mesophyll cytosol, is an important parameter that affects the estimate of *V*_pmax_. We have used *g*_m_ = 1 mol m^-2^ s^-1^ bar^-1^ here, together with the temperature dependence of *g*_m_ measured for *S. viridis* (Ubierna *et al*., 2017), however a variety of values have also been reported (Barbour *et al*., 2016; Ermakova *et al*., 2021b; Osborn *et al*., 2016; Pathare *et al*., 2022). Estimating mesophyll conductance in C_4_ species is difficult and requires measuring C^18^O^16^O isotope discrimination. If reliable measurements of *in vitro* maximum PEP carboxylase activity can be obtained, Equation 52 in von Caemmerer (2021) can be used to estimate *g*_m_. This approach was used by Ubierna *et al*. (2017). Unless *g*_m_ has been independently estimated, we recommend using the pre-set value of 1 mol m^-2^ s^-1^ bar^-1^.

The mitochondrial day respiration rate (*R*_d_) has been pre-set to 1 in the *A*/*C*_i_ parameters; this value is conventionally used in the absence of an experimentally determined estimate (von Caemmerer, 2021). For C_3_ species, the *R*_d_ and CO_2_ compensation point can be calculated as the *y*- and *x*-intercepts of the linear fit of *A* measurements at *C*_i_ values below 50 µbar; however, for C_4_ species this method often provides unrealistic values (Yin *et al*., 2011). As such, these values are not included as outputs of the tool. If a more accurate value of *R*_d_ is desired, we recommend either determining *R*_d_ from fitting *A*/*I* curves performed on the same leaf or constraining *R*_d_ to a set fraction of the leaf’s dark respiration rate (Sharkey, 2016).

### Fitting a light response curve

It is well recognized that the light response of C_4_ photosynthesis frequently does not saturate under ambient CO_2_ partial pressures (Cousins *et al*., 2006; Leakey *et al*., 2006). The shape of light response curve is determined by Equation 34 in von Caemmerer (2021) which, as for the C_3_ photosynthetic model, remains empirical. Importantly, the model considers the optimal partitioning of electron transport capacity between the C_4_ and C_3_ cycle, as discussed in detail in von Caemmerer and Furbank (1999) (also see: Peisker, 1988; von Caemmerer, 2000). Here the partitioning has been set at *x* = 0.4 (Table 1). The model now also better considers the contributions of *J*_cyc_ for accommodating the ATP demands of C_4_ photosynthesis. Using the model, *A*/*I* curves of C_4_ leaves can be described with the parameters *R_d_* and *J*_max_ (the maximum linear electron transport rate). The fitting of the *A*/*I* curve uses Equation 40 of von Caemmerer (2021).

To fit an *A*/*I* curve, one first updates leaf temperature, atmospheric pressure, O_2_ concentration and *α* in the spreadsheet (see details in ‘Fitting a CO_2_ response curve’ section). *A*, *C*_i_ and irradiance are then entered. Pressing the calculate button will solve for the empirical curvature factor (θ), *R*_d_ and *J*_max_ values which minimise the sum of squares between experimental and modelled data. After converging on a solution, the θ, *R*_d_ and *J*_max_ will be the output for the modelled electron transport limited CO_2_ assimilation rate at the measurement temperature. The maximum electron transport rates through *J*_cyc_ (*J*_max cyc_) and PSI (*J*_max 1_) are also determined from *J*_max_ and *f*_cyc_ (Equation 6). These electron transport rates are also calculated at 25°C and at the optimum temperature of electron transport (*T*_opt_) to facilitate reporting (Table 1). The quantum yield of CO_2_ assimilation (*ϕ*_CO2_) and light compensation point (*Γ*_light_; the irradiance at which *A* is zero) are calculated as the slope and *x*-intercept of the linear fit of measurements at irradiances below 200 µmol m^-2^ s^-1^.

### Important *A*/*I* fitting considerations

Along with CO_2_ assimilation rate, stomatal conductance also varies with irradiance, although at a much slower pace. This can lead to large variations in *C*_i_ when measuring light response curves. Incorporating *C*_i_ into the model calculations helps to improve the fitting routine. However, minimising variability in *C*_i_ over the course of an *A*/*I* measurement is preferable. To achieve this, some thought should be given to how quickly irradiance is changed to obtain a light response curve (McAusland *et al*., 2016).

While we advise measurements are performed on light-adapted leaves, it is important to consider whether light response curves are measured with increasing or decreasing irradiances, as this can impact parameter estimates. Measuring curves with increasing irradiances may result in lower CO_2_ assimilation rates due to deactivation of C_4_ enzymes at lower irradiances (Arce Cubas *et al*., 2022; Doncaster *et al*., 1989; Leegood and Furbank, 1984). Performing *A/I* curves with decreasing irradiance may help ensure enzyme activation but, in cases where plants are high light sensitive, slow relaxation of NPQ or photoinhibition may lead to lower *A* at lower irradiances, therefore resulting in underestimated *J*_max_ and *ϕ*_CO2_. While slow relaxation of NPQ could be mitigated by allowing more time at each irradiance step, photoinhibition typically results in long-term decrease of photosynthetic efficiency (Malnoë, 2018). Figure 4 provides examples of *A*/*I* measurements performed on *S. viridis* with increasing or decreasing irradiances. Fitting of the curve measured with increasing irradiance demonstrates θ values that are inconsistent between biological replicates and are lower than those previously reported, but *J*_max_ values that approach those expected (Massad *et al*., 2007; Sonawane *et al*., 2018). Comparatively, fitting of WT curves measured with decreasing irradiances provide θ values that are more consistent between biological replicates and are similar to those previously reported, but lower than expected *J*_max_ values (further discussed below) (Massad *et al*., 2007; Sonawane *et al*., 2018).

**Figure 3.**
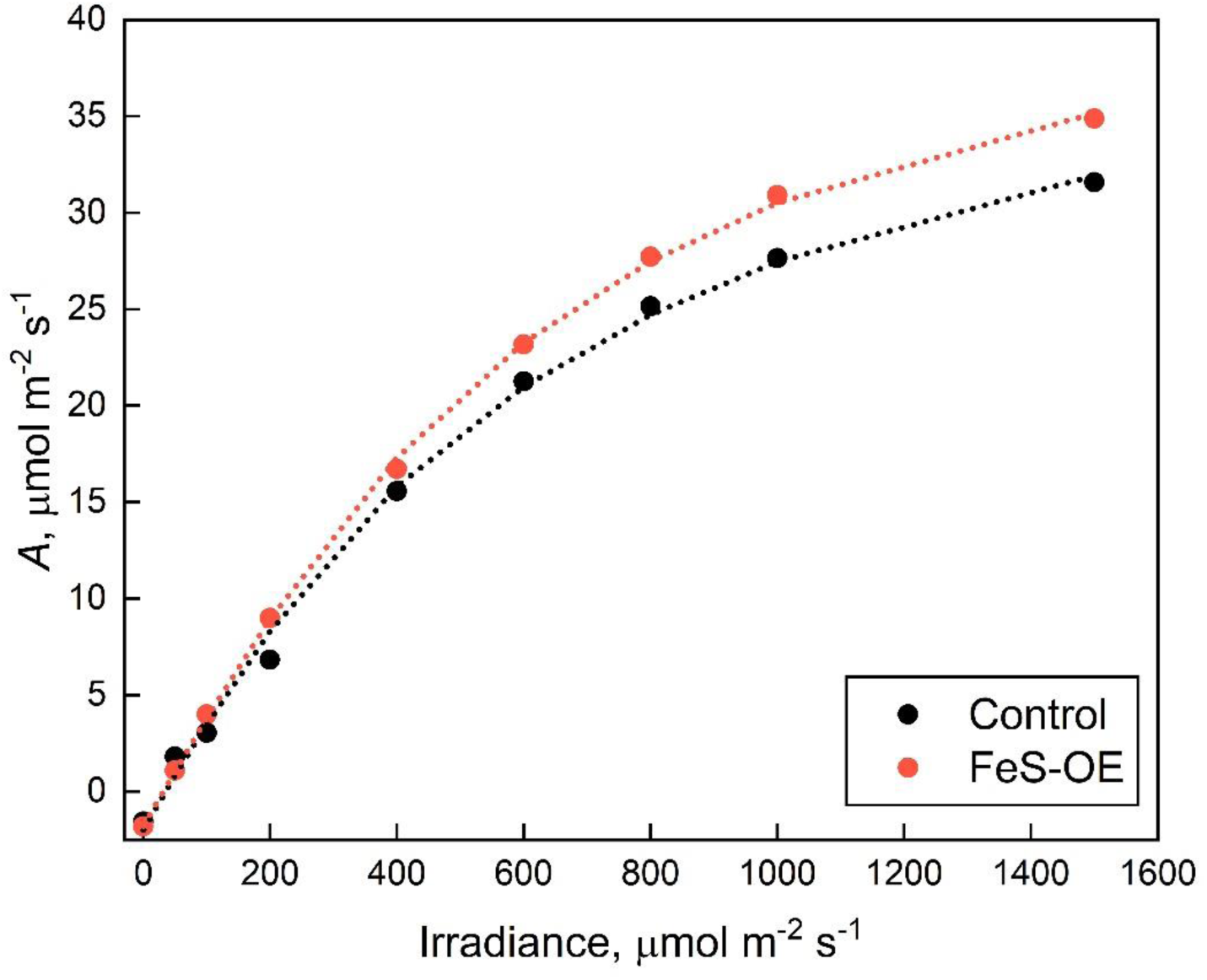
Fitting of the light response curves of CO_2_ assimilation (*A*) from Control and Rieske FeS overexpression (FeS-OE) *S. viridis* plants. Data originally reported in Ermakova *et al*. (2019). Dotted lines represent the fit of modelled data to the measured data points. The fitted values for FeS-OE plants are *R*_d_ = 1.79 µmol m^-2^ s^-1^, *J*_max_ = 182.89 µmol m^-2^ s^-1^, θ = 0.54, *ϕ*_CO2_ = 0.054 and *Γ*_light_ = 30.68 µmol m^-2^ s^-1^. For Control plants, *R*_d_ = 2.01 µmol m^-2^ s^-1^, *J*_max_ = 176.00 µmol m^-2^ s^-1^, θ = 0.37, *ϕ*_CO2_ = 0.040 and *Γ*_light_ = 25.91 µmol m^-2^ s^-1^.

**Figure 4.**
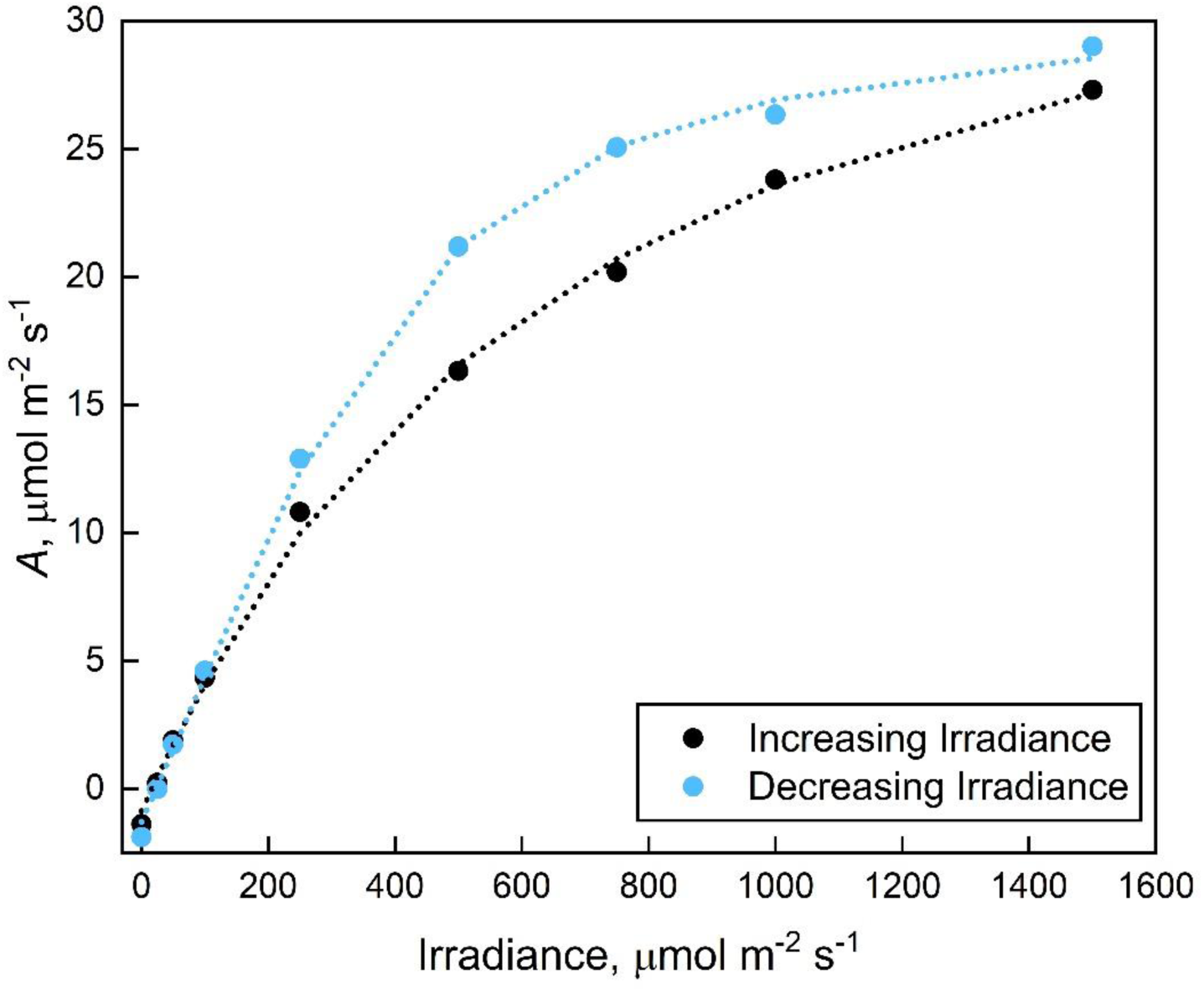
Fitting of light response curves of CO_2_ assimilation (*A*) measured with increasing and decreasing irradiances. Data from *S. viridis* leaves originally reported in Woodford *et al*. (2024). Dotted lines represent the fit of modelled data to the measured data points. For increasing irradiance, *R*_d_ = 0.88 µmol m^-2^ s^-1^, *J*_max_ = 152.25 µmol m^-2^ s^-1^, θ = 0.09, *ϕ*_CO2_ = 0.057 and *Γ*_light_ = 21.70 µmol m^-2^ s^-1^. For decreasing irradiance, *R*_d_ = 1.29 µmol m^-2^ s^-1^, *J*_max_ = 126.25 µmol m^-2^ s^-1^, θ = 0.83, *ϕ*_CO2_ = 0.065 and *Γ*_light_ = 26.72 µmol m^-2^ s^-1^.

As with *A*/*C*_i_ measurements, determining the maximum electron transport rate and disentangling electron transport and Rubisco limitations can be difficult. To resolve this problem, we recommend fitting *A*/*I* curves using measurements made under ambient or higher *C*_i_ at irradiances at which electron transport is expected to be limiting (i.e., <1500 µmol m^-2^ s^-1^). Using the fitting tool shows that *S. viridis* lines with Rieske FeS overexpression have increased *J*_max_ values, suggesting that an increase of assimilation seen in *A*/*C*_i_ at high *C*_i_ is due to increased electron transport capacity in the transgenic plants (Figures 2,3).

The *J*_max_ values reported by our tool are likely to be lower than previous literature values due to the suggested increase in *f*_cyc_ (Figure 5). To complement this change, we have also included a parameter for the rate of maximum electron transport through *J*_cyc_ which is determined from *J*_max_ and *f*_cyc_ using Equation 6. Following the derivations by von Caemmerer (2021), it is possible to sum *J*_max_ and *J*_max cyc_ to get a maximum electron transport rate through PSI (*J*_max 1_) (Figure 5b).

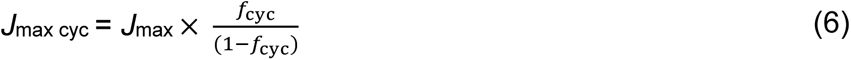

**Figure 5.**
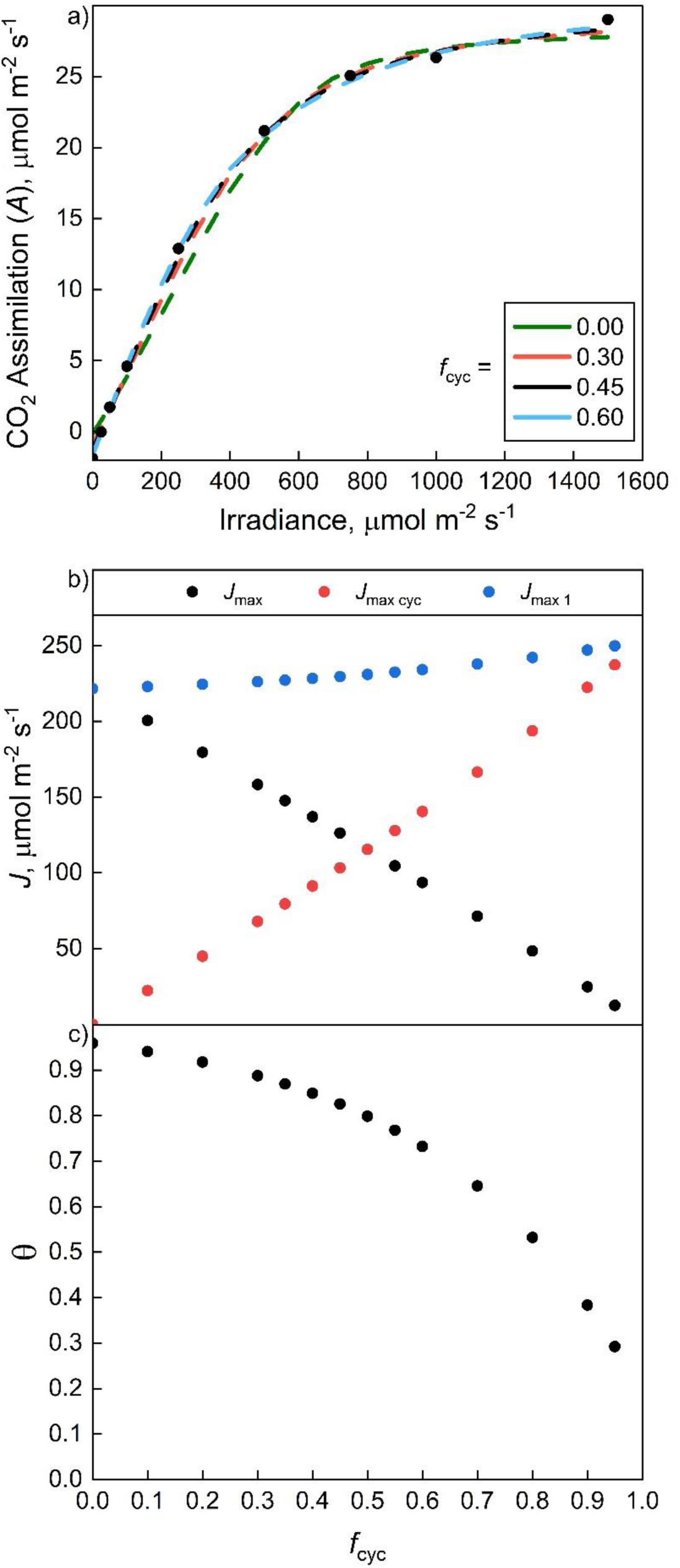
The proportion of electron transport occurring through cyclic electron flow (*f*_cyc_) influences the output parameters of light response curve (*A/I*) fitting. (**a**) Fitting *S. viridis A/I* curve measured with decreasing irradiance (from figure 4) with varying *f*_cyc_ values. (**b**) The output values for the maximum electron transport rates through linear electron flow (*J*_max_), cyclic electron flow (*J*_max cyc_), and Photosystem I (*J*_max 1_) obtained by fitting data from (a) with varying *f*_cyc_ values. (**c**) The output values for the curvature factor (θ) obtained by fitting data from (a) with varying *f*_cyc_ values.

The values for θ determined by our tool are influenced by the choice of *f*_cyc_ (Figure 5c). While θ values above 0.7 have been previously reported for both C_3_ and C_4_ *A*/*I* curves (Evans, 1989; Sonawane *et al*., 2018), the θ values derived by our tool may be lower than 0.7. This is because both θ and *f*_cyc_ act as curvature factors that influence the shape of the fit and are generally inversely proportional (Figure 5c). As a consequence, the potential interaction between θ and *f*_cyc_ should also be considered when interpreting outputs. It is possible that changes in θ could be reflective of changes in *J*_cyc_; however, the complexity of the relationship between θ and *f*_cyc_ requires further investigation. Importantly, the relationship between *J* and irradiance remains empirical and should not be over interpreted.

## Conclusion

The fitting of *A*/*C*_i_ and *A*/*I* curves to modelled data provides insight into the underlying biochemistry of C_4_ photosynthesis. By keeping fitting routines as simple as possible, our fitting tool provides streamlined, meaningful biochemical characterisation of *A*/*C*_i_ and *A*/*I* curves performed on C_4_ leaves. This should simplify reporting of the C_4_ gas exchange measurements and facilitate comparison between different experiments and species, allowing users to explore the variation in biochemistry of C_4_ photosynthesis. Combining *A*/*C*_i_ and *A*/*I* curves with estimates of Rubisco and Cytochrome *b*_6_*f* abundance and/or activity could help resolve limitations at high light and high *C*_i_. Fitted electron transport rate through PSII can be compared to estimates of electron transport rate from chlorophyll fluorescence, now frequently measured concurrently with gas exchange in many portable gas exchange systems.

## Supporting information

Supplemental File 1 - C4 Gas Exchange Fitting Tool

## Acknowledgements

This work was supported by a Research Scholarship from the Grains Research and Development Corporation to RW (UMO2501-001RSX) and the Australian Research Council’s Discovery Project to ME and RTF (DP230100175).

## Author Contributions

SvC, RW and ME designed the research; ME, SvC and RTF supervised research; RW updated model parameterisation and developed the Excel fitting tool; RW, SvC and ME wrote the paper; all authors discussed the results and contributed to the final manuscript.

